# Rt3DE-based finite element analysis of functional tricuspid regurgitation and RV free wall approximation

**DOI:** 10.64898/2026.07.13.736182

**Authors:** Davide Tondi, Simona Vailetta, Francesco Sturla, Riccardo Vismara, Emiliano Votta

## Abstract

**Purpose:** Functional tricuspid regurgitation (FTR) is driven by right ventricular (RV) remodeling, annular dilation, and papillary muscle dislocation. Free wall approximation (FWA) has been proposed to treat FTR by addressing RV dilation, but its effects on tricuspid valve (TV) biomechanics remain unclear. We present a real-time 3D echocardiographic (rt3DE)-based finite element framework to quantify TV biomechanics under FTR, and preliminarily apply it to assess FWA effects.

**Methods:** Subject-specific models were developed from rt3DE data of three dilated porcine hearts in an ex-vivo mock-loop. TV geometries at end-diastole and peak systole (PS) were complemented by parametric chordae tendineae and hyperelastic tissue properties. TV closure was simulated under a standard pressure load and image-based annular motion. After tuning chordae length to replicate the PS ground truth in FTR, FWA was simulated as 30% and 60% approximations along three anatomical directions (anterior-posterior, A-P; anterior-septal, A-S; anterior-septal wall, A-SW).

**Results:** In FTR simulations, median geometric errors ranged from 1.16 to 1.26 mm; median stress ranged from 56.4 to 74.7 kPa. FWA simulations predicted regurgitant orifice area (ROA) reductions by 53-99%, albeit overestimating the residual ROA vs. in vitro ground truth when starting from particularly extreme FTR conditions; concomitantly, a median stress reduction by 8-43% vs. FTR conditions was predicted.

**Conclusion:** Preliminary data suggest that our rt3DE-based framework can reliably quantify FTR-related TV biomechanics and that post-FWA biomechanics depends on initial FTR conditions. A larger cohort is required to verify the method and obtain statistically significant results.

## 1 Introduction

Tricuspid valve (TV) function arises from the coordinated interaction among its structural components: leaflets, annulus, chordae tendineae (CTs), and papillary muscles (PMs). Alterations in right ventricular (RV) loading conditions can induce RV pressure and/or volume overload, leading to chamber dilatation and tricuspid annulus (TA) enlargement. These morphological changes impair leaflet coaptation, resulting in functional tricuspid regurgitation (FTR) through two primary mechanisms: antero-posterior leaflet displacement associated with TA enlargement, and leaflet tethering driven by RV dilatation. FTR affects an estimated 1.6 million individuals in the United States alone [1] and is an increasingly recognized public health concern [2].

Current therapeutic approaches for FTR focus predominantly on the TA and, in some cases, directly target TV leaflets [3]; more recently, transcatheter tricuspid valve replacement has also become available [4]. However, these strategies largely neglect RV dilatation and the associated leaflet tethering, despite evidence demonstrating their crucial role in persistent or recurrent regurgitation. Indeed, when RV dilatation remains uncorrected, increased TV tenting volume has been identified as a key predictor of residual FTR following tricuspid annuloplasty [5]. Furthermore, progressive RV remodeling can continue postoperatively, predisposing patients to recurrent FTR at follow-up. Re-operation for recurrent FTR carries substantial perioperative and long-term mortality [6], underscoring the need for preventive and more effective treatment strategies.

Inspired by advances in the management of functional mitral regurgitation, subvalvular correction strategies that directly address RV dilatation have recently been proposed for FTR [7, 8]. Among these, free wall approximation (FWA) aims at approximating the RV free wall to the interventricular septum through a suture thus reducing the inter-papillary distance, thereby targeting the root cause of leaflet tethering rather than its downstream consequences at the annular or leaflet level. By counteracting PM displacement, FWA is expected to restore leaflet coaptation and, in the longer term, to promote favorable RV remodeling with secondary reduction of annular dimensions. Early clinical evidence from a small patient cohort suggests encouraging short-term outcomes, with no recurrence of FTR at one-year follow-up [9]. Nevertheless, the achievement of durable leaflet coaptation remains challenging [10], and several open questions persist regarding the optimal suture direction, the degree of approximation required, and the biomechanical impact of PM repositioning on valve function.

A key obstacle to answering these questions lies in the incomplete understanding of how the interplay between annular enlargement and subvalvular tethering driven by RV dilatation ultimately governs valve competence [11]. Because these factors are inherently coupled and patient-specific, their relative contributions to regurgitation, and consequently the optimal corrective strategy, cannot be reliably inferred from clinical imaging alone. These challenges highlight the need for a mechanistic framework capable of capturing the three-dimensional geometry and biomechanical interactions of the TV-RV complex.

Biomechanical analyses can be performed through high-fidelity finite element (FE) models based on clinical imaging, which capture patient specific geometry and in vivo motion of boundary regions. In particular, image-based FE models have been successfully applied to simulate mitral valve (MV) systolic biomechanics from multiple clinical imaging modalities [12–15], enabling detailed analyses of leaflet stress and strain [16], chordal loading, annular dynamics’ influence [17], and the effects of surgical and transcatheter interventions [18].

Building on that experience, FE modeling is increasingly being extended to study TV biomechanics in physiological conditions, in case of FTR, and following device implantation [19–23]. However, building a high-fidelity FE model is more challenging for the TV than for the MV: TV anatomy is more complex; its anterior anatomical position makes medical imaging more challenging and often limits image quality; reconstructing it from imaging data is further complicated by the thinness of TV tissues; and available data on the mechanical properties of TV tissues are still scarce.

On this basis, the present study has two objectives. First, advancing TV high-fidelity FE modeling leveraging on real-time 3D echocardiography (rt3DE). Second, preliminarily quantifying TV biomechanics in FTR conditions and following FWA.

To pursue both objectives within a controlled and reproducible setting, we relied on *in vitro* experiments performed on three porcine right hearts passively pulsating within a controlled mock loop. For each heart, FTR conditions were reproduced and then treated through FWA along different suture directions and shortening levels. For each condition, rt3DE was acquired along with direct endoscopic imaging. Rt3DE acquired in every tested configuration enabled i) the reconstruction of TV 3D geometry at end-diastole (ED) and peak systole (PS), ii) the FE simulation of TV systolic closure under FTR and post-FWA conditions, and iii) the validation of FE simulations.

## 2 Material and methods

The workflow implemented to obtain the relevant FE simulations for each TV is summarized in Fig. 1: rt3DE data acquired in FTR conditions were processed to reconstruct the 3D discretized geometry of annulus, leaflets, and PMs at ED and at PS. The reconstruction at ED was complemented with a paradigmatic set of CTs and used to feed the FE model of the valve. The reconstruction at PS was used to calibrate the stress-free length of CTs. Upon calibration, TV biomechanics during closure from ED to PS was quantified by forward FE simulations in FTR conditions and after six FWA variants encompassing three directions of the tethering suture and two levels of suture shortening.

**Fig. 1.**
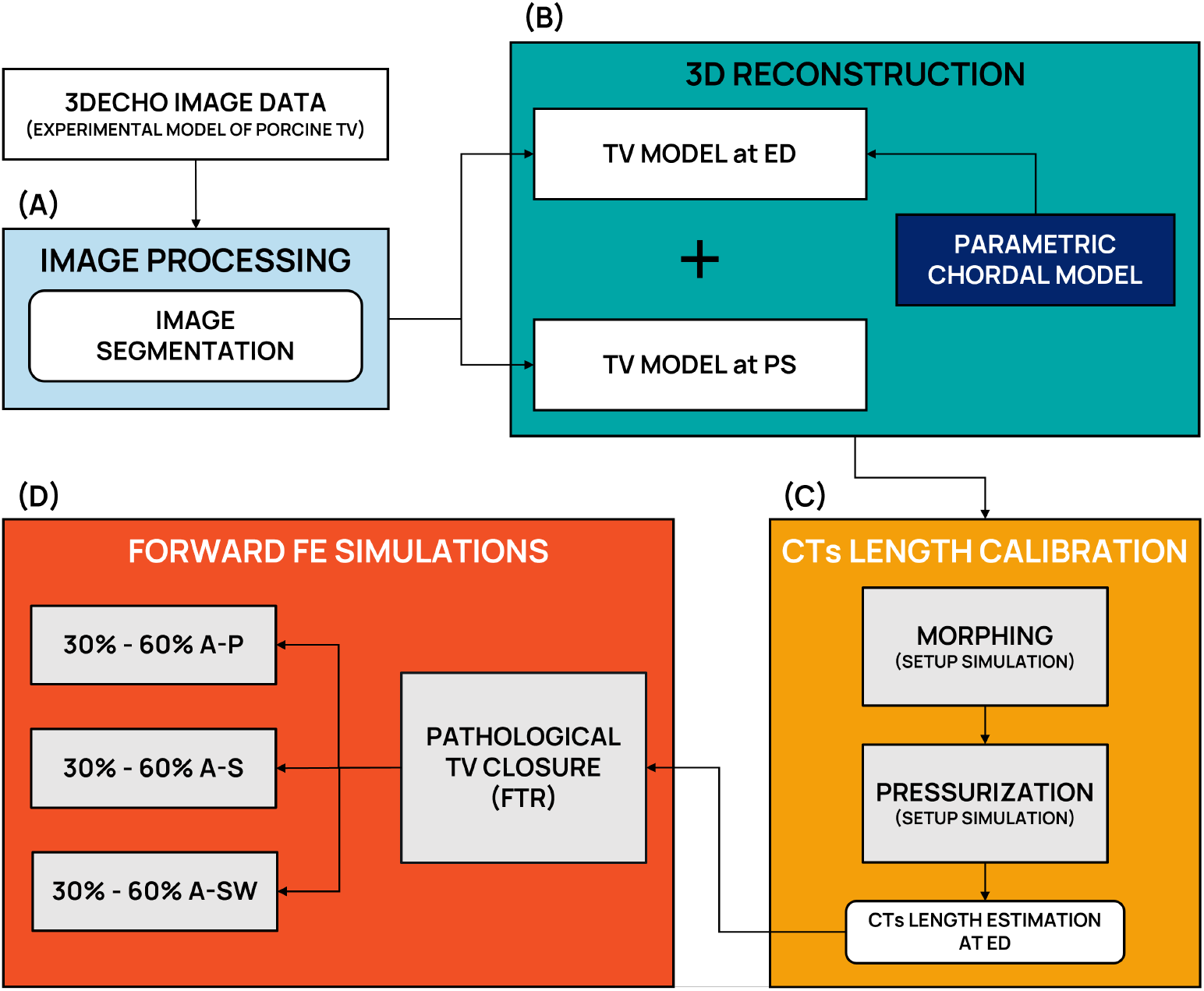
3D tricuspid valve (TV) modelling and simulation pipeline for functional tricuspid regurgitation (FTR) and free wall approximation (FWA)-based correction assessment. (A) 3D echocardiographic datasets acquired from ex-vivo porcine hearts were segmented using in-house algorithms to generate subject-specific TV geometries at end-diastole (ED) and peak-systole (PS). (B) At ED, the TV model was reconstructed including leaflet surfaces and a parametric representation of the chordae tendineae (CTs). (C) CT lengths were calibrated through two preliminary set-up simulations: “morphing”, to identify CT insertions on TV leaflets at PS, and “pressurization”, to identify CT tissue working point at PS and hence CT initial length. (D) Pathological valve behavior under FTR conditions was then simulated, followed by a series of FWA simulations aimed at evaluating the biomechanical impact of subvalvular correction. FWA simulations reproduced papillary muscle repositioning along three anatomical directions: anterior-posterior (A-P), anterior-septal (A-S), and anterior-septal wall (A-SW), with two levels of approximation (30% and 60% reduction of the initial inter-papillary distance) applied in each case. The predictive capability of the finite element computational model was assessed by comparing the resulting valve configurations across these scenarios. All simulations (light grey boxes) were performed through the finite element solver ABAQUS/Explicit 3DEXPERIENCE R2017x (SIMULIA, Dassault Systèmes).

### 2.1 Mock-loop design and data collection

Rt3DE was acquired on porcine right hearts in a previous *in vitro* study [24]. Briefly, porcine right hearts were integrated into a pulsatile mock loop equipped with a PC-controlled pump and an adjustable simulator of pulmonary impedence, which allowed to obtain controlled heart rate, time-dependent trans-pulmonary flow rate and RV pressure. Physiological resting conditions were set (60 bpm, 70 mL stroke volume). The pulmonary impedance was set to obtain a 15 mmHg mean pulmonary artery pressure under FTR conditions. These were obtained thanks to the tendency of porcine right hearts to dilate when loaded by physiological pressure levels, resulting in TA enlargement and leaflet malcoaptation due to excessive tethering, following the standardized protocol developed and validated by [25]. FWA was induced by passing a suture through the RV and adjusting its length via a 3D-printed mechanical winch, enabling the controllable simulation of FWA. FWA was performed in three anatomical directions aligned with papillary muscle positions, i.e., anterior-posterior (A-P), anterior-septal (A-S), and anterior-septal wall (A-SW) (Fig. 2.A). For each direction, the suture length was reduced by 30% and 60%, to simulate increasing degrees of approximation, through the winching mechanism (Fig. 2.B). In every experimental condition, TV atrial views were acquired through a fiberscope, and rt3DE was acquired using an ECG-gated Philips IE33 system equipped with an X7-2t probe, which was manually positioned on the epicardium. Acquisitions were imported into QLab (Philips Healthcare) and exported as Cartesian DICOM files.

**Fig. 2.**
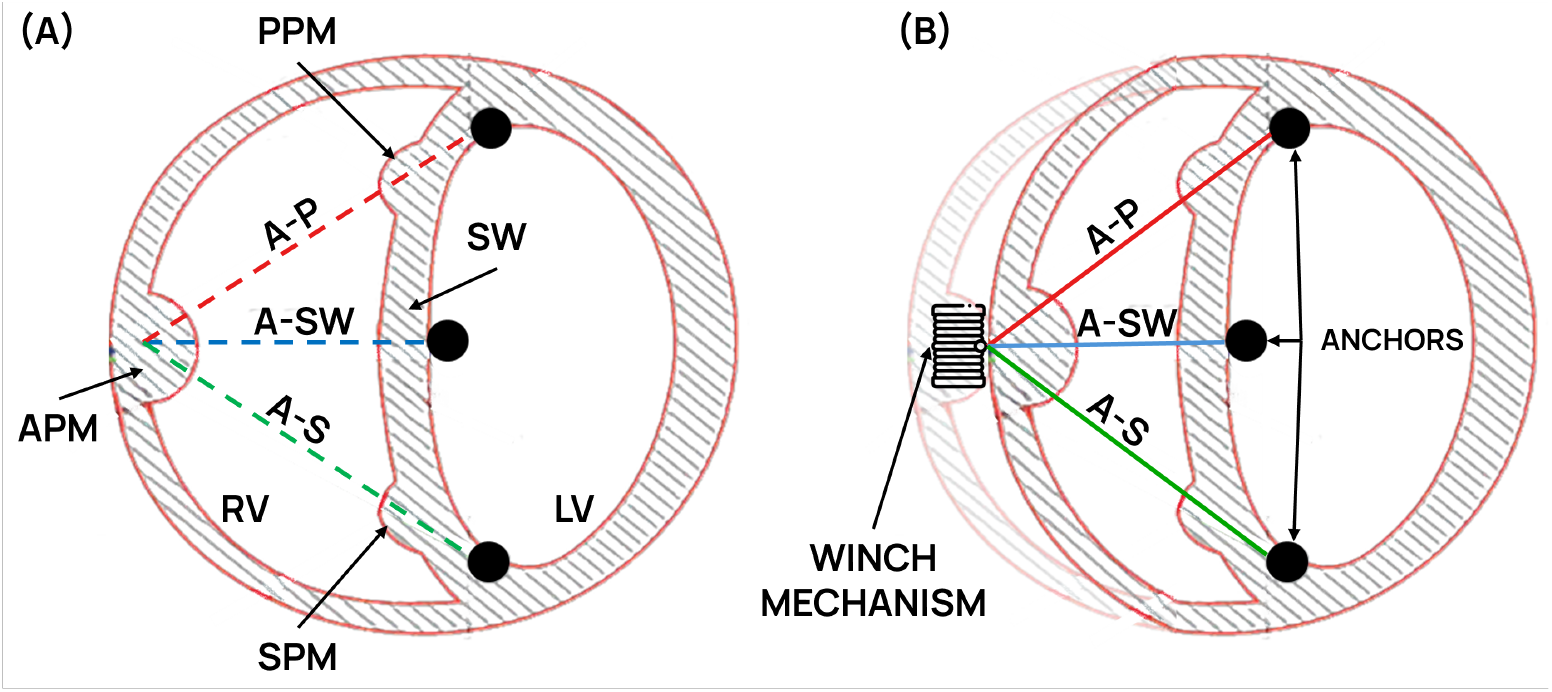
(A) The three anatomical directions, as seen in a short-axis section of the right ventricle (RV), considered for the free wall approximation (FWA): anterior-posterior (A-P), anterior-septal (A-S), and anterior-septal wall (A-SW). LV: left ventricle; PPM: posterior papillary muscle; APM: anterior papillary muscle; SPM: septal papillary muscle. (B) Schematic of the effects of suture shortening through a winching mechanism along one of the three approximation directions.

### 2.2 Image Segmentation

An *in-house* MATLAB script (MathWorks, Natick, Massachusetts, USA) was used to navigate the 3D ultrasound data, define a cylindrical reference frame with the *z*-axis centered and orthogonal to the TV orifice, and interpolate the volumetric data onto 18 radial planes evenly rotated around the *z*-axis.

Using a dedicated graphical user interface implemented in MATLAB [26], rt3DE data acquired in FTR conditions were processed to trace TV structures on each image plane at the time points corresponding to ED and PS (Fig. 1.A). All tracings were performed by a single operator with more than four years of experience in atrioventricular valve tracing across multiple imaging modalities. Borrowing the approach previously adopted for the MV [14], at the ED frame on each radial plane two points were identified on each leaflet at the annular insertion and at the free margin, and additional points were selected along the leaflet profile. For each leaflet, all points from the annular insertion to the free margin were interpolated using a cubic spline and subsequently resampled at 32 equally spaced locations.

At the PS frame, the same approach adopted at the ED frame was replicated on image planes where leaflets did not coapt, i.e., where the free margin of each leaflet was visible. On image planes where leaflets coapted, i.e., where it was not possible to identify the free margin of each leaflet, a single leaflet profile was traced, interpolated with a cubic spline, and resampled at 64 equally distributed positions.

For each frame and radial plane, the contours of the PM heads were also traced when visible.

### 2.3 Reconstruction of TV 3D geometry at ED

TV 3D geometry at ED was reconstructed following [27] (Fig. 1.B). For each point obtained by sampling tracings, the coordinates defined in the local 2D reference system of the echocardiographic image plane were transformed into the global 3D reference frame based on the information available in the Cartesian DICOM files.

#### 2.3.1 TV leaflets

As in [27], the 3D cartesian coordinates of the annular points were used to identify their centroid 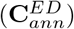 and least-square fitting plane 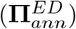, characterized by outward normal 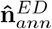. All the points generated by leaflet tracing and resampling were transformed into a cylindrical reference frame with the origin in 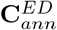 and the z-axis aligned with 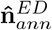 (Fig. 3.A), forming a point cloud organized in 32 curvilinear parallels and 36 curvilinear meridians, which were regularized using Fourier fitting functions and spline interpolating functions, respectively. Both Fourier and spline curves were then resampled at an approximate spacing of 0.4 mm along their respective directions. Since all simulations were performed through the finite element solver ABAQUS/Explicit R2017x (SIMULIA, Dassault Systèmes), the resulting leaflet surface (Ω^ED^) was automatically remeshed using Gmsh [28] with a prescribed element size of 0.4 mm, and discretized using three-node shell elements (ABAQUS S3R type) (Fig. 3.B).

**Fig. 3.**
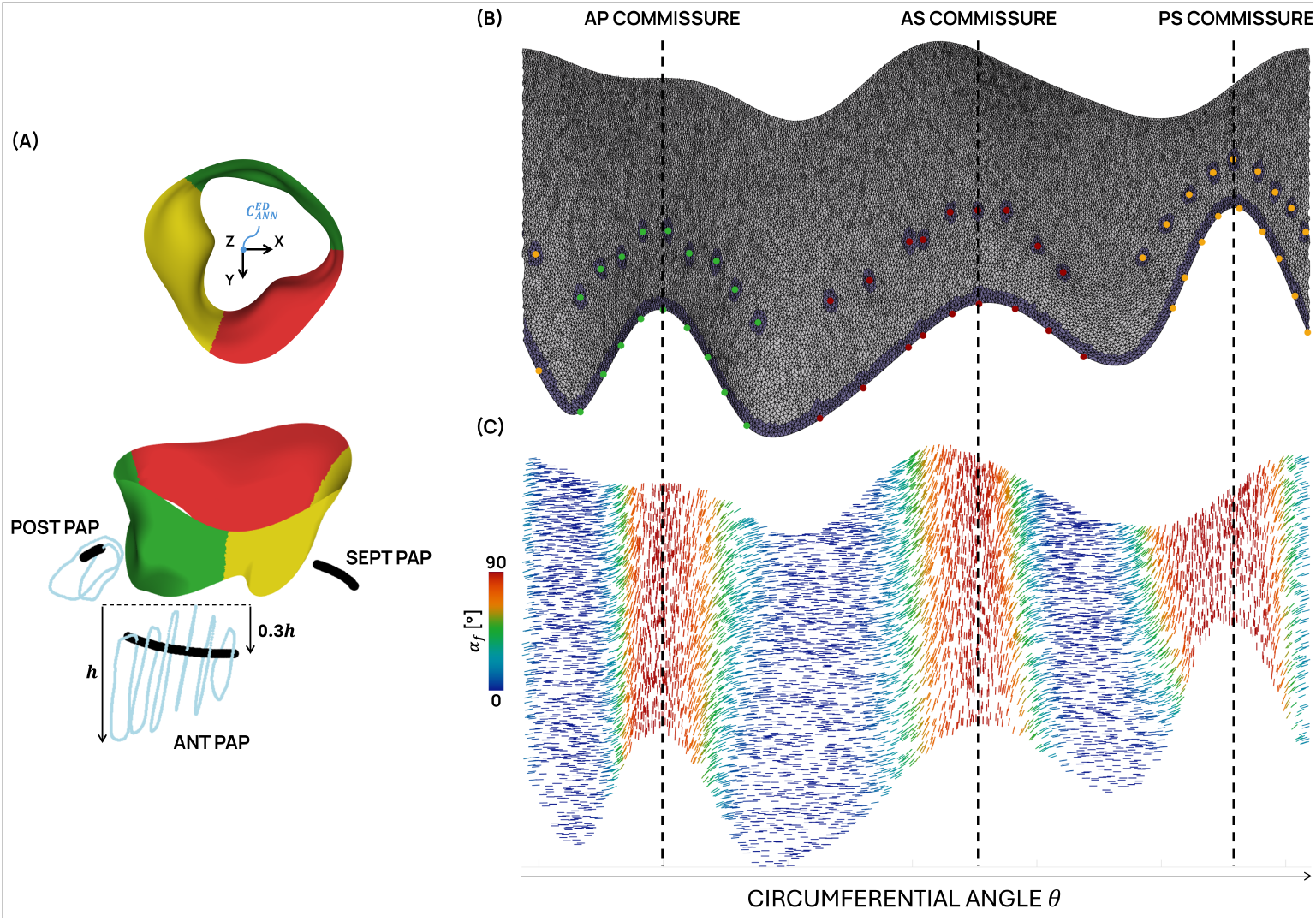
(A) Representative TV model shown from two perspectives: atrial view (up), with the local cylindrical reference system centered at the tricuspid annulus centroid 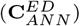, and lateral view (down), displaying the papillary muscle (PM) point clouds (light blue) derived from radial plane segmentations and the corresponding fitted circular arcs. Leaflets are color-coded: anterior (yellow), posterior (green), and septal (red). (B) Unfolded representation of the TV leaflets in the circumferential-longitudinal plane, showing the valve mesh with reinforcing elements (purple) at chordae tendineae (CT) insertion sites. Markers indicate CT attachment nodes, color-coded by papillary muscle of origin: green, anterior-posterior PM (18 CTs); yellow, postero-septal PM (20 CTs); red, antero-septal PM (18 CTs). (C) Prescribed fiber orientation angle *α*_f_ distribution across the unfolded valve surface. Fibers are predominantly circumferential (*α*_f_ = 0°) in the central region of each leaflet and rotate toward a longitudinal orientation (*α*_f_ = 90°) in the commissural regions, following a sigmoidal transition governed by Eq. (1). The horizontal axis reports the circumferential angle *θ* with respect to the tricuspid annulus axis.

Through the `*NODAL THICKNESS` keyword, shell elements were assigned a thickness field with larger values at the basal region, near the annulus, and at the free edge and a gradual decrease toward a minimum at the central region of each leaflet, in agreement with previous observations [29]. To this aim, the anterior, posterior, and septal leaflets were first identified on the triangulated surface by locating the commissural points. These were determined as the points of maximum curvature along the free margin curve, which delineate the boundaries between adjacent leaflets. Leaflets’ centroids were assigned minimum thickness values of 0.62 mm, 0.49 mm, and 0.57 mm, respectively, obtained by reducing by 10% the mean thickness values reported by Heyden et al. [30] for each leaflet. Nodes located along the free margin and annulus were assigned a uniform maximum thickness of 0.75 mm, derived from the highest mean leaflet thickness reported [30] for the anterior leaflet, further increased by 10%. These thickness values were interpolated by parametric sinusoidal functions, which were sampled at every node of the leaflet discretized geometry. This procedure resulted in average leaflet thicknesses across the three TV models of 0.66 mm for the anterior leaflet, 0.57 mm for the posterior leaflet, and 0.62 mm for the septal leaflet.

The mechanical behavior of TV leaflet tissue was described using the Holzapfel-Gasser-Ogden (HGO) constitutive model [31], assuming a single family of collagen fibers. Fiber orientation (*α*_f_) was defined using a parametric description with respect to a local cylindrical coordinate system centered on the axis passing through the valvular orifice (Fig. 3.C). In accordance with experimental observations by Cochran et al. [32] on human MV leaflets, fibers were assumed to be predominantly circumferential in the central region of each leaflet (*α*_f_ = 0°) and predominantly longitudinal in the commissural regions (*α*_f_ = 90°). The fiber orientation angle was assumed constant along the radial direction of the leaflet surface and allowed to vary only circumferentially. In the transition region between the central and commissural zones, fiber orientation was prescribed using a sigmoidal function, *α*_f_ defined as

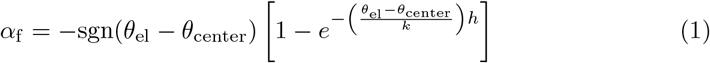

In this formulation, *θ*_el_ denotes the angular coordinate of the element centroid, *θ*_center_ identifies the angular position of the leaflet midline, and *s* = *k*(*θ*_center_ *− θ*_comm_) defines the characteristic angular scale of the transition. The sgn function ensures a symmetric distribution of fiber rotation with respect to *θ*_center_, enforcing a right-hand-rule convention such that fibers rotate progressively from the leaflet center toward the commissures. The parameters *h* and *k* control the steepness and extent of the sigmoidal transition and were set to *h* = 4 and *k* = 0.6 to reproduce the angular extent of the circumferential and longitudinal plateau regions reported by [32]. Specifically, the central circumferential plateau (*α*_f_ = 0°) spanned 20% of the total leaflet angular extent, while the longitudinal plateau (*α*_f_ ≈ 90°) occupied the outer 10% on each side in the commissural regions. The constitutive parameters of the model, *c*_10_, *k*_1_, *k*_2_, and *κ*, were identified for each leaflet by fitting equi-biaxial tensile test data acquired by Amini et al. [33] on porcine TV leaflet samples (Table 1).

**Table 1.** Leaflet-specific constitutive parameters of the HGO model used to describe the mechanical behavior of the TV leaflets, identified for each leaflet by fitting equi-biaxial tensile test data reported by [33] on porcine TV leaflet samples.

| | $c_{10}$ [MPa] | $k_1$ [MPa] | $k_2$ [-] | $\kappa$ [-] |
| --- | --- | --- | --- | --- |
| Anterior | 2.91E-03 | 828.60E-03 | 119.25 | 3.08E-01 |
| Posterior | 1.00E-03 | 282.93E-03 | 45.25 | 2.93E-01 |
| Septal | 1.86E-03 | 379.52E-03 | 49.20 | 3.08E-01 |

Reinforcing elements were introduced at CT insertions into the leaflets (Fig. 3.B). For each leaflet node corresponding to a CT insertion, the adjacent leaflet elements were duplicated to create an additional local layer, whose thickness was set to one tenth of the average thickness of the surrounding leaflet elements. These reinforcing elements were assigned an isotropic, linear elastic material. The Young modulus was set equal to the secant elastic modulus of CT tissue at a nominal strain of 0.07, corresponding to a value of 10 MPa, while the Poisson ratio was set to 0.475. The reinforcing elements were meant to imitate the presence of a transition in mechanical properties from CT to leaflet tissue, and to prevent from unrealistic mesh distortions and stress artifacts associated with the concentrated load transfer at CT insertions.

#### 2.3.2 Papillary muscles

PMs visible in the image volume were reconstructed from segmentations obtained on radial planes and subsequently expressed in the local cylindrical reference system. For each PM, the head was modeled as a circular arc fitted to the point cloud of the PM segmentation, with an angular extent matching that of the segmented PM and laying on a plane parallel to the best-fit plane of the TA.

The arc was positioned at 30% of the PM height (*h*) along the local axial (*z*) direction of the cylindrical reference system, where 0% corresponds to the PM point closest to the annular plane (Fig. 3.A). This choice was made to approximate the effective region of chordal attachment while ensuring geometric consistency across models.

If a PM was not visible within the imaging volume, its head was modeled as a circular arc positioned below the corresponding commissure, with both the angular extent and the axial distance from the annular best-fit plane set equal to the mean values of the corresponding quantities measured for the remaining reconstructed PMs.

The whole process resulted in a set of CT origins 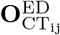 for the *j*-th PM head, where *i* indicates the origin of the *i*-th CT.

#### 2.3.3 Chordae Tendineae

CTs were modelled as non-branched strings and discretized into linear truss elements (ABAQUS, T3D2 type elements) with a characteristic length of 2 mm and a constant cross-sectional area of 0.17 mm^2^ [34]. The distribution of CT insertions on TV leaflets was borrowed from [20], based on the experimental observations therein reported: 18 CTs stem from both the anterior and septal PMs, and 20 CTs stem from the posterior PM. For each PM, CTs were evenly distributed between marginal chordae, inserted on the leaflet free edge, and second-order chordae, inserted at approximately two thirds of the leaflet height. For each CT, the origin on the corresponding PM arc was defined as the arc point with the minimum Euclidean distance to the leaflet insertion point of that chorda.

The mechanical behavior of CTs was described using the Weiss constitutive model [35], assuming that CT tissue is incompressible and reinforced by a single family of collagen fibers aligned with the CT axis. The total strain energy density function *W* is expressed as the sum of an isotropic contribution from the ground substance matrix (*W*_ISO_) and an anisotropic contribution from the collagen fibers (*W*_f_):

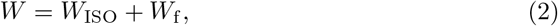

Under the assumption of incompressibility, *W*_ISO_ is given by

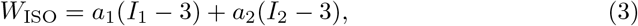

with *I*_1_ and *I*_2_ being the first and second invariants of the right Cauchy-Green deformation tensor **C**. *W*_f_ is a function of the stretch ratio of the material along the fiber direction (*λ*_f_), such that:

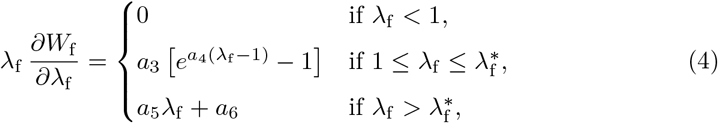

where 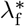 denotes the stretch ratio at which fibers become fully recruited, marking the transition from the exponential toe region to the linear regime.

The Cauchy stress (***σ***) is correspondingly decomposed as

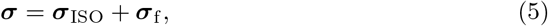

where the isotropic contribution (***σ***_ISO_) is

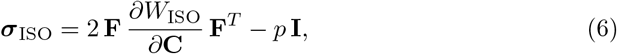

with *p* being the Lagrangian multiplier introduced to cope with the incompressibility constraint, and the fiber contribution (***σ***_f_) is

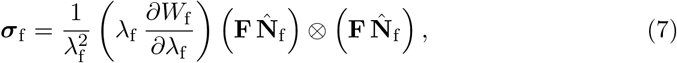

where 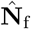 is the unit vector defining the initial fiber direction in the reference configuration.

The constitutive parameters *a*_*i*_, with *i* = 1, 6, and 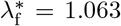 were identified by fitting the model to the experimental stress-strain data derived from uniaxial tensile tests reported by [34] (Table 2). The Weiss model was implemented in a user-defined material ABAQUS subroutine (VUMAT).

**Table 2.** Constitutive parameters were obtained by fitting experimental stress-strain data derived from uniaxial tensile tests [34].

| $a_1$ [MPa] | $a_2$ [MPa] | $a_3$ [MPa] | $a_4$ [-] | $a_5$ [MPa] | $a_6$ [MPa] | $\lambda_f^*$ [-] |
| --- | --- | --- | --- | --- | --- | --- |
| 16.852 | -1.651 | 16.159 | -5.681 | -57.580 | 56.580 | 1.063 |

### 2.4 Reconstruction of TV 3D geometry at PS

#### 2.4.1 TV leaflets

The TV leaflet surface at PS (Ω^PS^) was reconstructed from the points obtained by tracings performed at the PS frame. As for Ω^ED^, prior to surface reconstruction the coordinates of these points were transformed from the image reference frame into the cylindrical reference frame previously defined at ED (Section 2.3.1).

Depending on the degree of FTR-related leaflet hypomobility, a different reconstruction strategy was adopted at PS.

In absence of leaflet coaptation, Ω^PS^ was reconstructed following the same procedure adopted for the ED configuration: leaflet points were transformed into the local cylindrical reference frame and organized into a structured point cloud of curvilinear parallels and meridians, regularized using Fourier fitting functions and spline interpolating functions, and subsequently triangulated and discretized using S3R elements. In case of partial leaflet coaptation, Ω^PS^ was reconstructed through a dedicated approach: the 576 leaflet points were clustered into 16 concentric profiles, interpolated using NURBS functions and automatically triangulated through a Delaunay-based procedure. The resulting surface was smoothed using a Laplacian smoothing algorithm with a relaxation factor of 0.01, and finally modeled as a rigid shell surface discretized with S3R elements.

#### 2.4.2 Papillary muscles

The region of PM heads corresponding to CT origins were reconstructed at PS through the same approach adopted at ED (Section 2.3.2) to define the position of CT origins at PS. These were used to define the 3D displacement of each CT origin from ED to PS.

### 2.5 CTs length calibration

Since CTs are not directly visualized by standard clinical imaging modalities [36], a dedicated strategy was implemented to model CT lengths while reproducing ground-truth ultrasound-derived tracings. As in [27], prior to simulating valve closure, CT stress-free length at ED was calibrated through a three-step process:

1. *Morphing simulation* Ω^ED^ was driven to approximate Ω^PS^, yielding an estimated closed-valve configuration 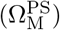. CT tissue was modeled as linear elastic and highly extensible, with an elastic modulus of 5 kPa, in order to avoid unrealistic CT motion during the morphing simulation while minimally affecting the local kinematics of Ω^ED^. A pressure load linearly increasing up to 5 mmHg was applied on the ventricular side of the leaflets to promote contact between Ω^ED^ and Ω^PS^, while annular nodal displacements between ED and PS were prescribed from rt3DE tracings and PM tips were kept fixed in their PS position. Based on the position of the nodes representing CT insertions on 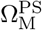, the loaded length of each CT at PS 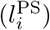 was estimated as the distance between the corresponding insertion node and the corresponding origin node on the relevant PM head.
2. *Pressurization simulation* CTs were modeled as nearly inextensible linear elastic elements with an elastic modulus of 100 GPa and a Poisson ratio of 0.475, consistent with the assumption of full collagen fiber recruitment along the chordal direction. A physiological pressure increasing up to 30 mmHg was applied to the ventricular surface of 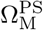. The corresponding CT axial stresses (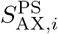 where *i* indicates the *i*-th CT) were computed.
3. *Final CT length tuning* based on 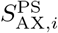 the axial stretch ratio of each CT (*λ*_AX,*i*_) was obtained by inverting the CT constitutive law using the computed axial stress. The stress-free CT length at ED was then calculated as 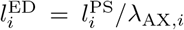. The new reference configuration of each CT was set by keeping its original orientation in the 3D space and the original position of its insertion on TV leaflets updated accordingly; a new position of its origin 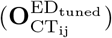 was thus defined.

### 2.6 Forward FE simulations

Following the calibration of CT initial length TV systolic closure from ED to PS was simulated under FTR and post-FWA conditions.

#### 2.6.1 Simulation of TV closure under FTR conditions

A two-step simulation was performed using Ω_ED_ as initial configuration of leaflets and annulus and 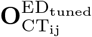 as initial position of CT origins. In step 1, which lasted 0.5 seconds, nodal displacements computed as:

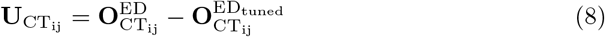

were imposed to the nodes representing the CT origins, while the nodes of annulus and leaflets were prevented from any translation.

In step 2, lasting 0.235s, a physiological time-dependent transvalvular pressure, increasing up to 30 mmHg, was applied on the ventricular side of Ω_ED_. All contact interactions, including leaflet self-contact, were modeled through a penalty contact algorithm with a friction coefficient of 0.05 [26]. Contextually, nodal displacements were prescribed to the nodes on the annular profile and to CT origins to replicate annulus and PM motion between ED and PS as obtained from rt3DE. Displacements of annular nodes were prescribed by enforcing a kinematically consistent deformation of each inter-commissural segment while preserving the original nodal discretization, such that the number of annular nodes within each segment remained unchanged throughout deformation. TV leaflet and CT tissues were modelled as described in Section 2.3.

#### 2.6.2 FWA procedure simulation

For each TV, each FWA variant, i.e., each combination of suture direction and shortening (Section 2.1), was simulated by a two-step FE forward simulation using again Ω_ED_ as initial configuration of leaflets and annulus and 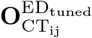 as initial position of CT origins.

In step 1, lasting 0.5s, the relevant post-FWA ED configuration 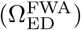 was obtained by:

- displacing CT origins to their post-FWA end-diastolic position 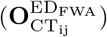. These positions were not reconstucted from post-FWA rt3DE, 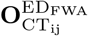 was computed as:

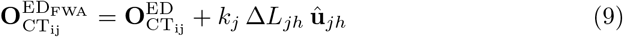

where *j, h* ∈ *{S, A, P, SW}* denote the PM or septal wall at each end of the suture, **û**_*jh*_ is the unit vector along the suture direction, and ∈*L*_*jh*_ is the prescribed suture shortening (Fig. 4.A). The scaling factor *k*_*j*_ ∈ [0, 1] governs the fraction of the total shortening assigned to each PM and was chosen to reflect the asymmetric mechanical response of the RV walls: for A-P and A-S directions, *k*_*A*_ = 0.8 and *k*_*P*_ = 0.2 or *k*_*S*_ = 0.2; for the A-SW direction, *k*_*A*_ = 0.8 and *k*_*S*_ = *k*_*P*_ = 0.1. This asymmetric distribution was motivated by the greater structural thickness [37, 38] and dominant load-bearing role [39] of the septal wall relative to the RV free wall, despite experimental evidence of lower myocardial stiffness in the septum [40];
- deforming the annular profile into its post-FWA end-diastolic configuration, as obtained directly from post-FWA rt3DE segmentation. Nodal displacements were defined following the same rationale reported in Section 2.6.1 (Fig. 4.B).

**Fig. 4.**
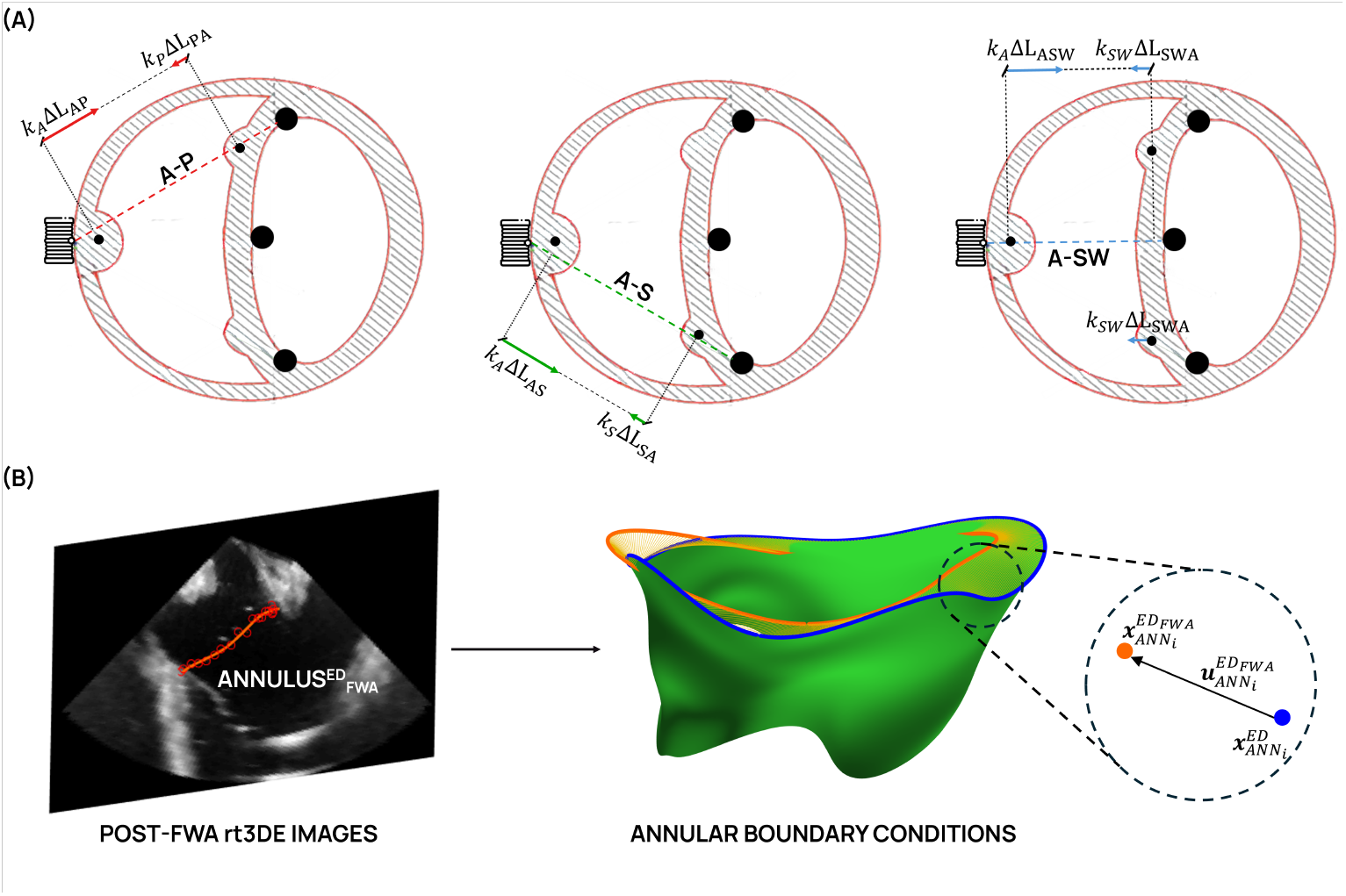
(A) Boundary conditions applied to CT origin nodes during step 1 of the FWA simulation, represented on a short-axis section of the right ventricle. The three anatomical suture directions considered for the free wall approximation (FWA) are shown: anterior-posterior (A-P), anterior-septal (A-S), and anterior-septal wall (A-SW). The scaling factors *k*_*j*_ governing the fraction of suture shortening assigned to each PM are reported for each direction. (B) Annular boundary conditions applied during step 1 of the FWA simulation. The post-FWA end-diastolic (ED) annular profile 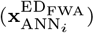 was obtained from segmentation of post-FWA rt3DE volumes. Nodal displacement vectors 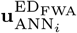 were computed as the difference between 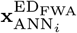 and the pre-operative ED nodal coordinates 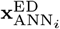, and prescribed to the annular nodes.

In step 2, lasting 0.235s, TV closure was simulated starting from 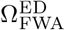. A time-varying transvalvular pressure load was applied on the ventricular side of the valve, following a systolic pressure profile and reaching a peak value of 30 mmHg. Concurrently, annular deformation between ED and PS, as derived from tracings on post-FWA rt3DE, was simulated by imposing displacements to the annular nodes, and PM motion from ED to PS was simulated assuming it equal to the one imposed in FTR simulations (Section 2.6.1).

#### 2.6.3 Effect of FWA-induced changes in annular kinematics

Ex vivo data highlighted a change in annular geometry and motion from ED to PS induced by FWA (Table 4). To investigate the relative contribution of the post-FWA changes in annular ED geometry and systolic motion to post-FWA TV biomechanics, an auxiliary set of simulations was performed. For each of the three TVs, the AP30 and AP60 configurations were re-simulated by maintaining the baseline, i.e., FTR-related, annular ED geometry and systolic motion (Section 2.6.1). Differences in the predicted leaflet stresses vs. the standard FWA simulations were quantified. Results of this analysis are reported in the Supplementary Material (Section S1).

## 3 Results

### 3.1 Baseline FTR simulations

#### 3.1.1 Geometric accuracy of predicted TV configuration at PS

For each considered TV, we computed the distribution of nodewise Euclidean distance (*ε*_D_) between the simulated closed TV configuration at PS, obtained from the TV closure forward simulation, and (Ω^PS^) (Fig. 5.A).

**Fig. 5.**
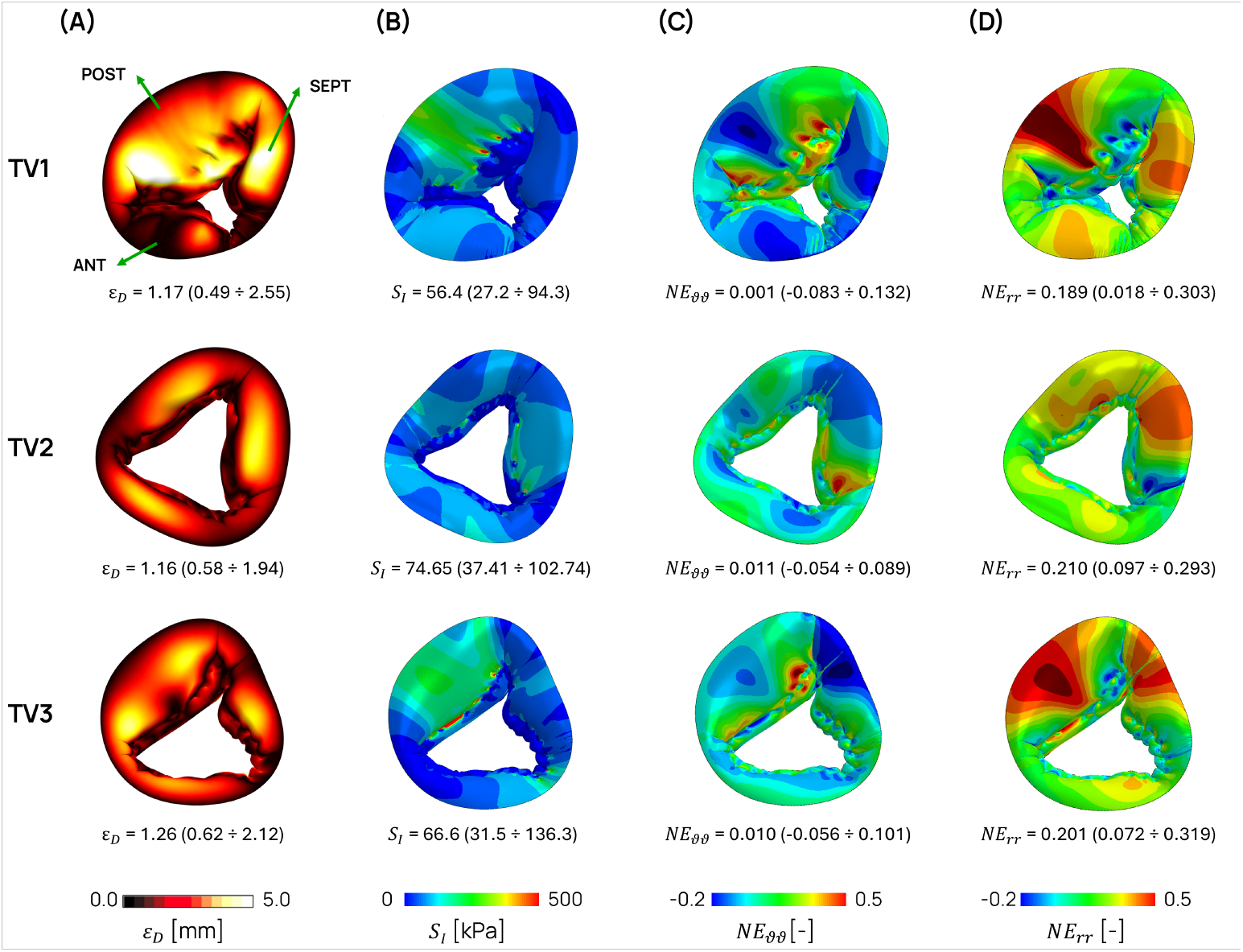
Peak-systolic geometric accuracy and biomechanical response of the tricuspid valve under functional tricuspid regurgitation (FTR) conditions for the three baseline configurations. (A) Atrial view of the node-to-node error, computed as the Euclidean distance (*ε*_D_, [mm]) between the simulated leaflet configuration at peak systole and the ground-truth leaflet geometry reconstructed from image segmentation. (B) Distribution of the maximum principal stress on the leaflet surface at peak systole (*S*_I_, [kPa]). (C) Distribution of the circumferential Green-Lagrange strain component (NE_*θθ*_). (D) Distribution of the radial Green-Lagrange strain component (NE_*rr*_). For all quantities, values are reported for each valve as the median with interquartile range (25^th^-75^th^ percentiles) computed over the entire leaflet surface.

Across all models, the median value of *ε*_D_ ranged from 1.16 mm in model TV2 to 1.26 mm in model TV3, against a voxel spacing of rt3DE imaging of 0.63 *±* 0.05 mm (mean *±* standard deviation of the in-plane and through-plane resolutions). Leaflet-specific analysis revealed greater variability: while for valves TV2 and TV3 balanced error distributions were obtained across all leaflets, for valve TV1 the posterior leaflet was characterized by markedly greater errors (median value of 1.98 mm) as compared to the anterior and septal leaflets. In general, larger *ε*_D_ values were observed in the central regions of the most extended leaflets, particularly the posterior leaflet in several models: in these regions, the simulated configuration was characterized by excessive bulging due to the reduced mechanical support by CTs. These evidences underscore the influence of leaflet geometry and subvalvular support on local simulation accuracy. The simulated PS configuration was further validated by qualitative comparison against video frames acquired with an endoscopic camera during in vitro testing on the mock-loop platform (Fig. 6). For all the three valves, the simulated FTR configuration well replicated the shape and extent of the regurgitant area observed in vitro: the anterior (yellow), posterior (green), and septal (red) leaflets exhibited coaptation patterns consistent with the endoscopic images, supporting the ability of the FE framework to capture the main features of TV closure under regurgitant conditions. Only for valve TV3 relevant discrepancies in regurgitant area shape were observed: the posterior leaflet was more mobile in the FE simulation than in the in vitro experiment, likely due to its particularly large extent that resulted in reduced chordal support and increased bulging. Also, the anterior leaflet appeared slightly underextended, possibly due to the underestimation of its annulus-to-free edge extent in the manual segmentation of rt3DE images.

**Fig. 6.**
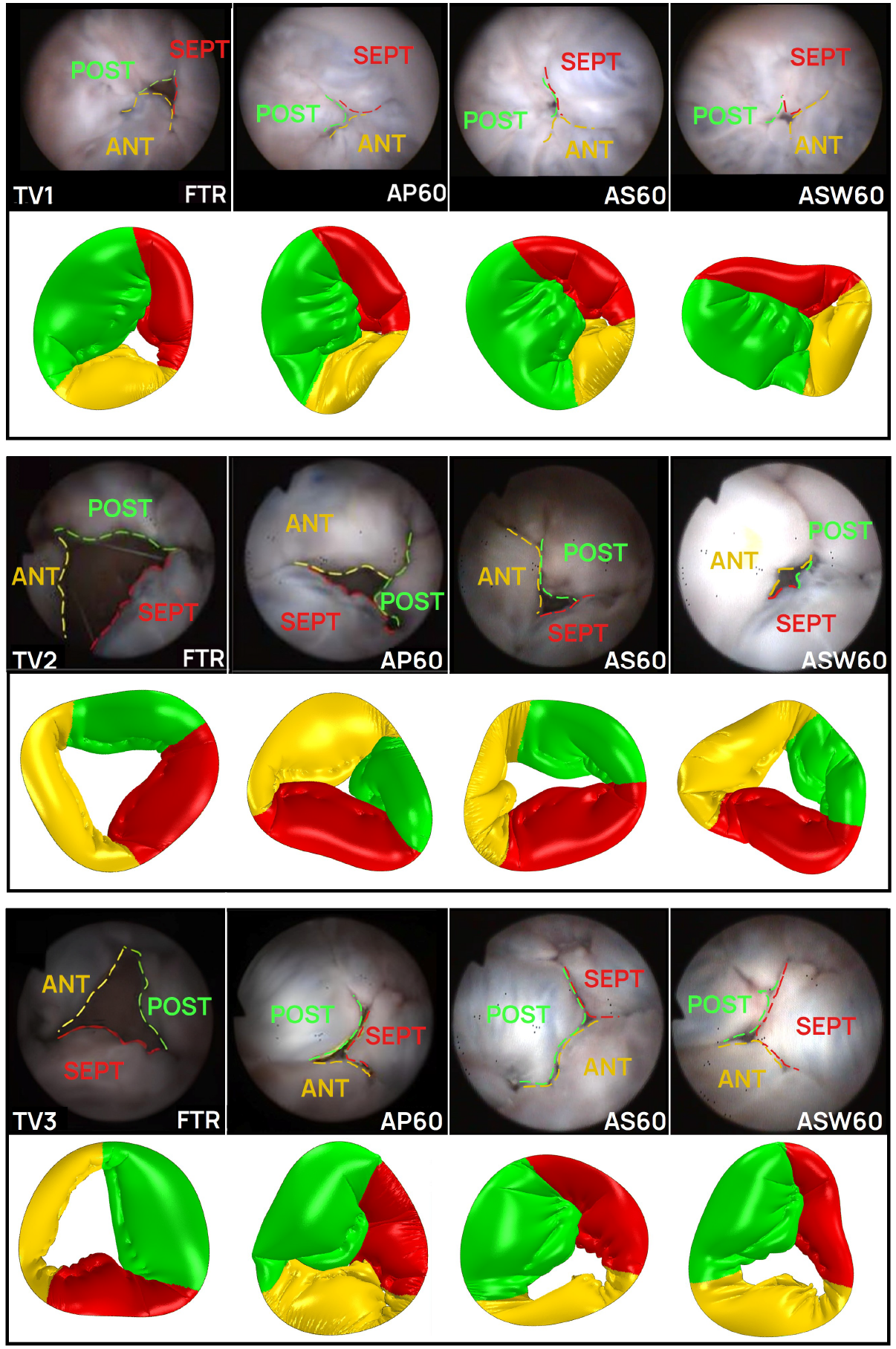
Qualitative comparison between peak-systolic tricuspid valve configurations from endoscopic video frames acquired during in vitro testing (upper row) and corresponding FE simulations (lower row) for three representative specimens (TV1, TV2, and TV3). Configurations are shown under functional tricuspid regurgitation (FTR) and after free wall approximation at 60% inter-papillary distance reduction along the three considered anatomical directions: anterior-posterior (AP60), anterior-septal (AS60), and anterior-septal wall (ASW60). In the simulated configurations, leaflets are color-coded: anterior (yellow), posterior (green), and septal (red).

#### 3.1.2 TV stresses and strains at peak systole

TV leaflet maximum principal stresses *S*_I_ were different among the three simulated TVs: the median *S*_I_ value ranged from 56.4 kPa in TV1 to 74.7 kPa in TV2. Moreover, while TV2 exhibited a fairly homogeneous stress field over the three leaflets, with leaflet-specific *S*_I_ median values of 71.6-77.8 kPa, in TV1 and TV3 the anterior and posterior leaflets were more stressed than the septal leaflet. For instance, in TV1 median *S*_I_ values of 73.7 kPa, 74.4 kPa, and 41.8 kPa were computed on the anterior, posterior, and septal leaflet, respectively (Fig. 5.B).

Leaflets experienced limited circumferential strains (NE_*θθ*_), with median values close to zero (from 0.001 in TV1 to 0.011 in TV2) (Fig. 5.C).

Radial strains (NE_*rr*_) were predominantly positive (Fig. 5.D), with median NE_*rr*_ ranging from 0.189 (TV1) to 0.210 (TV2). The posterior leaflet generally exhibited the highest NE_*rr*_ values, with peak values of 0.60 localized in the posterior and septal leaflet belly regions.

### 3.2 Post-FWA TV biomechanics

#### 3.2.1 Geometric Accuracy

As for simulated FTR conditions, the leaflet configuration at PS predicted by FWA simulations was compared against the corresponding ground-truth leaflet surfaces segmented from post-intervention rt3DE images at PS 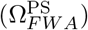 by computing *ε*_D_ (Fig. 7). For each valve, all post-FWA scenarios exhibited higher median *ε*_D_ values than the corresponding FTR condition. Within this general trend, inaccuracies in the prediction of leaflets configuration at PS depended on FWA direction and magnitude. When simulating FWA in the A-P direction (AP30 and AP60), a 30% approximation produced the largest *ε*_D_ median values in TV1 (1.70 mm) and TV3 (1.62 mm), against the values obtained when simulating FTR. In case of 60% approximation, *ε*_D_ median values decreased slightly but still exceeded those obtained when simulating FTR. These results were confirmed by the comparison against endoscopic video frames acquired during in vitro FWA testing (Fig. 6): although FE simulations captured the progressive reduction in regurgitant area induced by 30% and 60% A-P approximation, the shape of the computed regurgitant area did not always match the in vitro ground truth, with higher mismatches noted in TV2 for the AP60 configuration. When simulating FWA in the A-S direction (AS30 and AS60), errors were different across TVs. At 30% FWA, median *ε*_D_ values were moderately increased with respect to FTR, reaching 1.63, 1.32, and 1.39 mm for TV1, TV2, and TV3, respectively. However, in case of AS60, TV2 and TV3 exhibited larger discrepancies (by approximately 1.8 mm), while TV1 showed closer agreement, with errors closer to those observed in the FTR baseline.

**Fig. 7.**
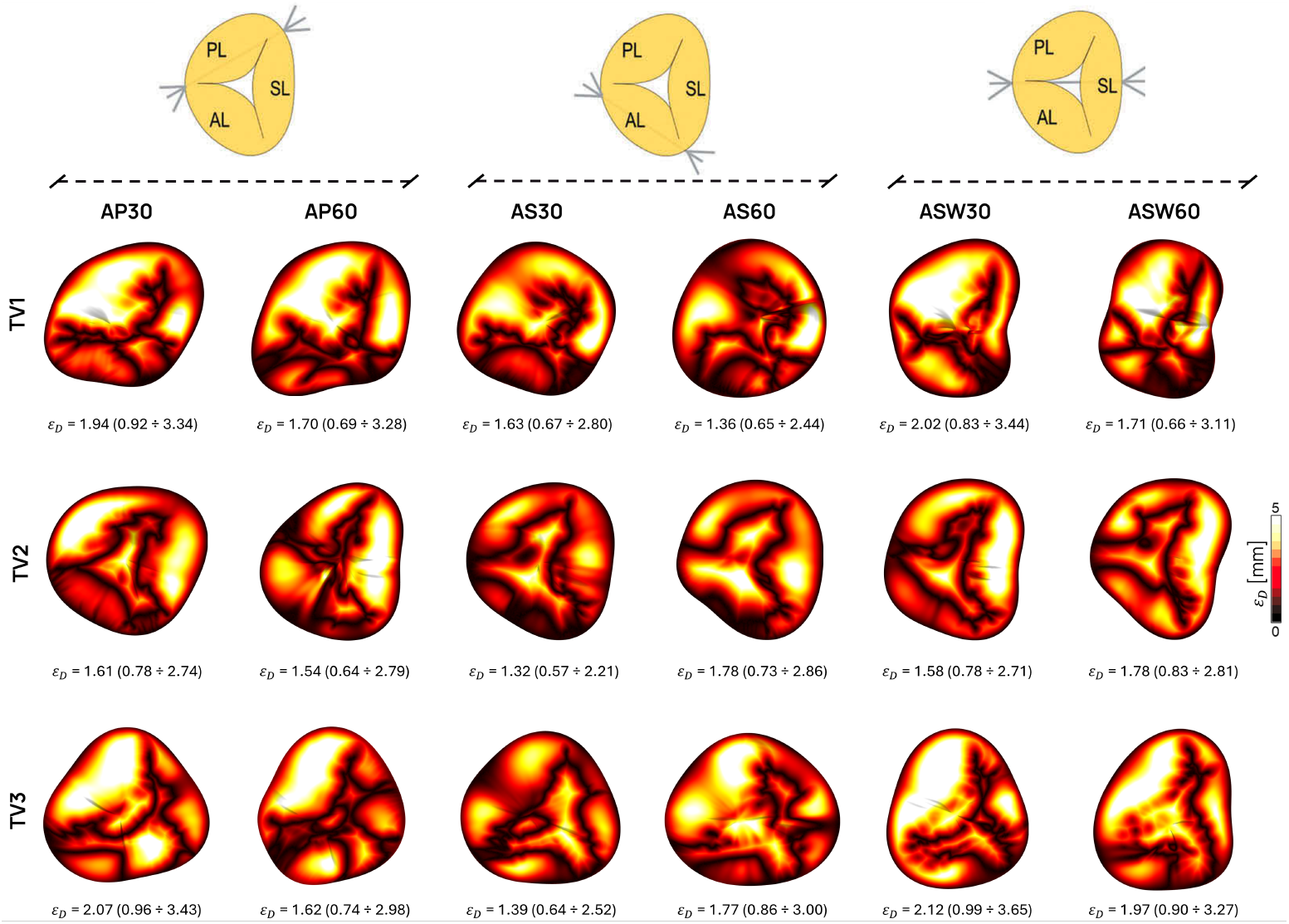
Peak-systolic geometric accuracy of the tricuspid valve under free wall approximation (FWA) conditions for the three analyzed samples. Nodewise Euclidean distance errors (*ε*_D_) were computed between the simulated closed TV configuration and the ground-truth leaflet surface reconstructed from rt3DE segmentation. FWA configurations include anterior-posterior (A-P), anterior-septal (A-S), and anterior-septal wall (A-SW) papillary muscle approximations at 30% and 60% inter-papillary distance reduction levels. Values are reported for each valve as the median with interquartile range (25^th^-75^th^ percentiles) computed over the entire leaflet surface.

When simulating FWA in the A-SW direction (ASW30 and ASW60), the highest *ε*_D_ values were obtained, particularly in TV3 where the median *ε*_D_ values was 2.12 mm and 1.97 mmm when simulating ASW30 and ASW60, respectively. Similarly, TV2 showed median errors of 1.58 mm and 1.78 mm.

#### 3.2.2 Leaflet stress distribution

According to FE simulations, FWA produced direction- and length reduction-dependent modifications in leaflet biomechanics and coaptation efficacy. The heterogeneous stress patterns observed under FTR conditions, with elevated *S*_I_ on the anterior and posterior leaflets and peak concentrations at chordal insertions, were variably attenuated depending on the approximation strategy (Fig.8). Concurrently, all FWA configurations reduced the regurgitant orifice area (ROA) relative to FTR baseline, though to a different extent depending on the TV (Table 3). To rule-out the effects of TV size on ROA assessment, we also computed %ROA as the ratio between ROA and the annular area of the corresponding FTR configuration, both projected on the annular best-fit plane.

**Table 3.**
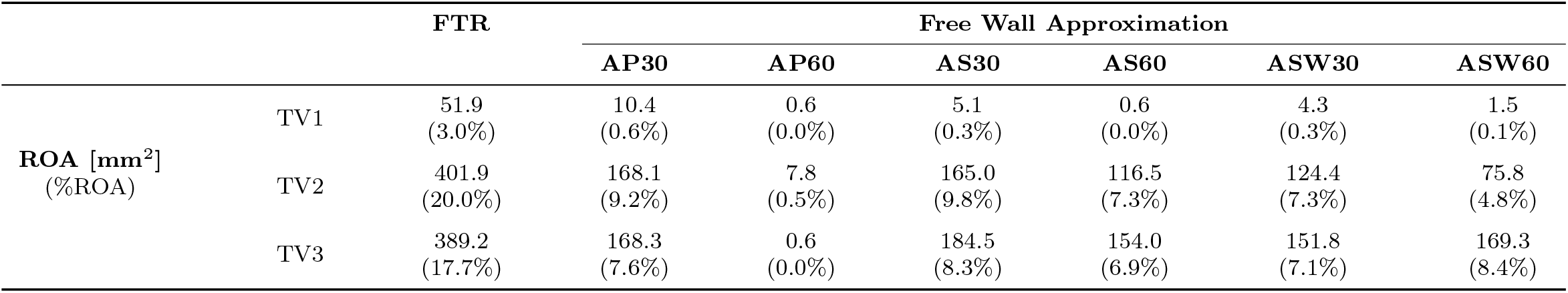
Regurgitant orifice area (ROA) at peak systole under functional tricuspid regurgitation (FTR) and free wall approximation (FWA) conditions. ROA was computed from the simulated systolic TV configuration. Percentage regurgitant area (%ROA) is expressed as the ratio of ROA to the 2D annular orifice area. FWA configurations include anterior-posterior (A-P), anterior-septal (A-S), and anterior-septal wall (A-SW) papillary muscle approximations at 30% and 60% inter-papillary distance reduction levels.

In case of FWA in the A-P direction, for a 30% approximation modest overall reductions in *S*_I_ were observed in TV1 and TV3 (*−* 17.7% and *−* 13.5%, respectively), though leaflet-specific responses varied substantially: in TV2, posterior leaflet stresses increased by over 50%, while anterior leaflet stresses decreased by nearly 40%. ROA was markedly reduced in TV1, while TV2 and TV3 retained significant residual orifice areas. At 60% approximation, *S*_I_ reductions became more pronounced (ranging from *−*22.7% to *−*36.6%), and near-complete coaptation was achieved across all specimens, with ROA approaching zero (%ROA ≤ 0.4%).

A-S approximation demonstrated variable global effects on leaflet stresses. For AS30 configurations, median *S*_I_ reductions ranged from *−*18.3% to *−*27.0%, though the posterior leaflet experienced higher stress in TV2 and TV3, with the latter increasing by +27.5% relative to FTR. ROA reductions at this level were comparable to A-P 30% 20 for TV1 and TV2, while TV3 retained a slightly larger orifice. At 60% approximation, stress responses diverged across specimens: TV3 showed negligible change in overall median *S*_I_ despite a *−*30.8% reduction in posterior leaflet stress. Notably, A-S 60% yielded less complete coaptation than A-P 60%, with residual %ROA above 5% in TV2 and above 7% in TV3

A-SW approximation yielded the most consistent *S*_I_ reductions across specimens. At 30% approximation, median stress decreased by 24-37% across models, with further reductions for ASW60 (by up to *−*43.4%). The symmetric displacement distribution (10% each at SPM and PPM) resulted in more balanced stress distributions as compared to asymmetric strategies. ROA reductions for ASW30 were comparable to those obtained when simulating other FWA directions (TV1: 4.3 mm^2^; TV2: 124.4 mm^2^; TV3: 151.8 mm^2^). At 60% approximation, A-SW further reduced ROA in TV2 (75.8 mm^2^, %ROA = 3.8%), though TV3 showed a minor increase relative to the ASW30 approximation (169.3 mm^2^, %ROA = 8.1%), complemented by increased anterior leaflet stresses (up to almost 2 MPa).

In all post-FWA simulated scenarios, peak leaflet stresses remained consistently localized at the insertions of CTs originating from the APM, which experienced the greatest displacement in PM repositioning. Elevated stresses propagated from these insertion sites toward the annulus, creating band-shaped regions of increased stress aligned with the direction of PM repositioning.

## 4 Discussion

While FE modeling of atrioventricular valves has been extensively developed for the MV [41], TV models remain comparatively underdeveloped owing to TV complex geometry and thinner leaflets, which complicate the accurate reconstruction of TV 3D geometry from clinical imaging, and to the limited availability of data on the mechanical properties of TV tissues [42]. In this study, we presented an ultrasound-based finite element modeling framework for quantifying TV biomechanics directly from rt3DE data acquired in three porcine hearts on a in-vitro mock loop [24]. The study brings three main methodological contributions to TV computational modeling: (i) the tuning of the chordal apparatus, which is not visible in standard clinical imaging, to reproduce with high-fidelity the TV systolic closure in a baseline scenario; (ii) the development of an automatic and reproducible workflow for simulating pathological FTR conditions; and (iii) the evaluation of the framework’s ability to predict the effects of FWA on TV systolic closure.

### 4.1 Geometric accuracy

A central challenge in image-based valve modeling is reconstructing the subvalvular apparatus, since standard imaging does not resolve the CTs [36]. Building on chordal calibration concepts developed for the MV [16, 27], we adapted this approach to the TV to closely replicate TV leaflet configuration at PS as reconstructed from rt3DE. Median distances between the simulated configuration and the rt3DE-based ground truth ranged from 1.16 mm to 1.26 mm across models, compared to a voxel spacing of approximately 0.63 mm; these values fall within the range of variability inherent in manual 3D echocardiographic segmentation: Nguyen et al. [43] reported inter-user intra-class correlation coefficient (ICC) values of 0.83 on average for TV annular measurements, and as low as 0.60 for mean lateral through-plane displacements. The largest discrepancies were typically localized in the belly region of the most extended leaflet, i.e., in the most extended leaflet regions characterized by reduced chordal support. This effect is inherent to the anatomically-informed chordal distribution adopted in this study, wherein CTs are mostly inserted on the leaflet free margin. An alternative approach based on uniformly distributed CTs [44], including attachments in the belly region, would have improved the agreement between the simulated and the ground-truth leaflet configuration in the baseline scenario, but would have compromised the prediction of post-FWA configurations. In uniformly distributed CT models, chords are also inserted in regions, such as near the annulus, where CTs are not present in the real valve; the initial length of each chord is then calibrated so that the model reproduces a target closed configuration in a given scenario without generating stress artifacts at the insertions. When simulating FWA, however, the PMs are repositioned and their distance from the CT insertions changes, and may increase relative to the baseline. As a result, chords whose length was optimal in the baseline scenario can become overly stretched to span the new PM-to-insertion distance, producing unrealistic stress concentrations and limiting the ability to predict post-FWA TV closure. Discrepancies between the simulated leaflet configuration at PS and the rt3DE ground truth increased when predicting the post-FWA scenarios, yet remained within 2-3 times the voxel spacing. Qualitative comparison against endoscopic video frames (Fig. 6) provided a more nuanced picture: for TV1, i.e., the valve characterized by the least severe malcoaptation and by the mildest annular dilation in FTR conditions, the simulated systolic configuration well reproduced the in vitro observations across all FWA directions. Instead, for TV2 and TV3, which were characterized by extreme annular dilation and malcoaptation, agreement between the computed post-FWA systolic configuration and the corresponding experimental evidence from endoscopic video frames was obtained only for when simulating FWA in the A-P direction, namely for the AP60 configuration. The AS60 and ASW60 configurations predicted by the simulations were less consistent with the endoscopic frames and overestimated the residual regurgitant area. It is reasonable to speculate that these discrepancies are related to the simplifying assumptions adopted to prescribe PM repositioning (Section 2.6.2) to reproduce the effects of FWA. Our results suggest the hypotesized direction of PM repositioning and the split of the reduction in suture length over the two PMs allows to reproduce the in vitro scenario when considering FWA in the A-P direction, but not when the A-S and A-SW directions are considered in severely regurgitant valves. It is also worth noting that, unlike the FTR validation, where both annular and PM positions were derived from segmentation, FWA simulations prescribed PM positions based not on image tracings but on the hypothesized pattern depending on the considered FWA direction, while annular geometry was matched to the segmented post-FWA profile. This methodological difference may also contribute to the increased geometric discrepancies observed in FWA configurations relative to FTR.

### 4.2 FTR biomechanics

Under FTR conditions, stress distributions exhibited marked inter-leaflet heterogeneity, with consistently higher values on the anterior and posterior leaflets as compared to the septal leaflet. This pattern is consistent with the greater mechanical demand placed on the mural leaflets: annular dilatation and PM dislocation associated to FTR preferentially increase their tethering [45]. Although the present work is not aimed at predicting leaflet remodeling, it could be of interest to investigate whether the leaflet regions exhibiting elevated stress levels in our models, extending from CT insertions toward the corresponding annular insertions, may correspond to regions more susceptible to progressive tissue remodeling. Experimental studies have reported that TV leaflets in FTR undergo adaptive thickening and stiffening [46], and that leaflet thickness may predict residual TR after repair [47].

The dominant role of radial elongation, with median NE_*rr*_ substantially exceeding circumferential strains, reflects the preferential compliance of TV leaflet tissue in the radial direction as the primary mechanism for accommodating closure under pathological loading [48].

### 4.3 Biomechanical effects of FWA

Although the very limited number of simulated valves and the hypotheses underlying the simulation of FWA require caution in interpreting computational results, our preliminary data suggest some potentially interesting trends. Importantly, the extent to which these trends reflect the true post-FWA behavior varied across configurations: as detailed below and in Section 4.1, the simulations reproduced the in vitro endoscopic observations (Fig. 6) for the A-P direction in all valves and for all directions in TV1, i.e., the least regurgitant valve, but only partially for the A-S and A-SW directions in TV2 and TV3, i.e., the two valves starting from extreme FTR conditions.

To facilitate the clinical interpretation of the simulated conditions, ROA values can be translated to the current TR grading scheme based on effective regurgitant orifice area (EROA) thresholds [49]: under FTR conditions, TV1 corresponds approximately to a moderate-to-severe TR grade, whereas TV2 and TV3 represent TR severities exceeding the torrential grade. It should be noted, however, that the anatomical ROA obtained from the computational model is not directly equivalent to the clinically measured EROA. From a fluid-dynamic perspective, EROA corresponds to the effective jet area at the vena contracta and therefore depends on both the geometry of the orifice and the local flow characteristics, and is generally smaller than the corresponding anatomical ROA [50]. The reported correspondence should thus be interpreted as indicative rather than as a direct quantitative match.

The first trend concerns the reduction in ROA vs. FTR conditions: across all considered FWA directions and approximation levels, simulations consistently predicted major ROA reductions, by 48%-99%, which were complemented by reductions in median leaflet *S*_I_ ranging from approximately 8% to 43% depending on direction and approximation level. The only exception was AS60 in TV3, where the overall median stress remained essentially unchanged. These effects depended on direction and magnitude of FWA-induced PM displacement, as well as on the annulus size and malcoaptation severity in FTR conditions. These outcomes suggest two remarks: extreme malcoaptation and annular dilation in FTR conditions may be associated to more complex FWA effects in terms of PM repositioning; this is well exemplified by valves TV2 and TV3, characterized by approximately eightfold ROA values as compared to TV1 (Table 3) and by larger annular dimensions (Table 4), whose post-FWA simulations likely did not correctly capture the real PM repositioning leading to the overestimation of the residual ROA. Moreover, the role of leaflet coaptation in limiting leaflet stresses is confirmed: whenever simulations overestimated the residual post-FWA malcoaptation, reductions in leaflet stresses vs. FTR conditions were markedly less evident.

**Table 4.**
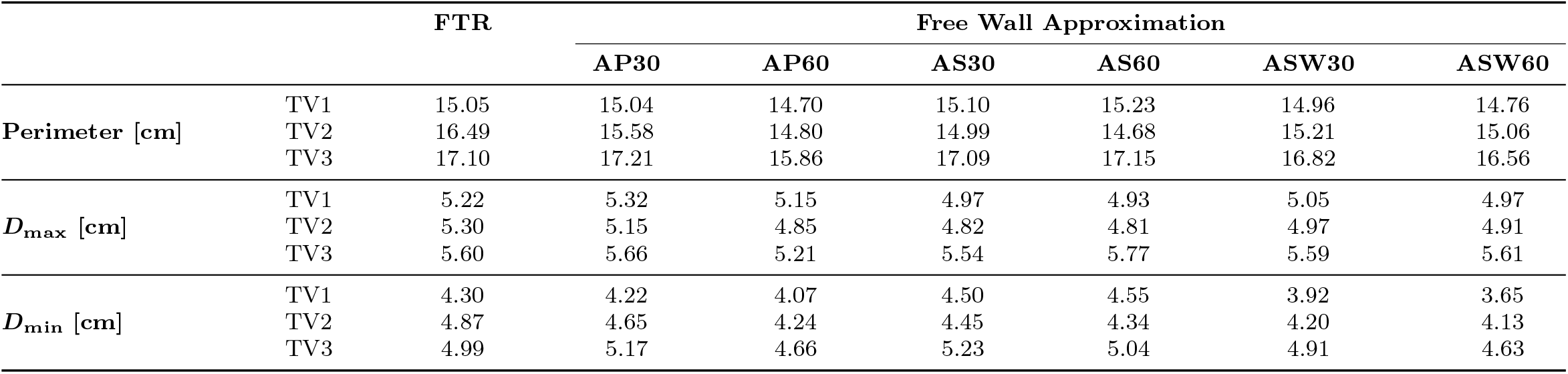
Morphological measurements of the tricuspid annulus at peak systole under functional tricuspid regurgitation (FTR) and free wall approximation conditions. Measurements were automatically derived from the ground-truth leaflet surfaces reconstructed from manual segmentation of rt3DE data acquired during in vitro testing. FWA configurations include anterior-posterior (A-P), anterior-septal (A-S), and anterior-septal wall (A-SW) papillary muscle approximations at 30% and 60% inter-papillary distance reduction levels. *D*_max_ = maximum annular diameter; *D*_min_ = minimum annular diameter.

Second, despite the overestimation of residual ROA for valves TV2 and TV3 when simulating A-S and A-SW approximation directions, simulations highlighted that in general ROA was progressively reduced when considering increasing approximation levels, no matter the approximation direction.

Third, the observed trend in ROA reduction vs. approximation level was paralleled by a similar trend in leaflet stresses at PS (Fig. 8): in general, post-FWA computed configurations were characterized by lower stress levels vs. the corresponding FTR configurations, and, for a given approximation direction, greater approximation levels led to a further reduction in leaflet stresses. The only exceptions to this general trend consisted in the configurations AS60 and ASW60 for valve TV3, wherein stresses increased vs. configuration AS30 and ASW30, respectively. Of note, in both these specific cases, the higher stress levels were localized in the AP commissure, whose free edge is a site of multiple insertions of CTs originating from the underlying anterior PM. When simulating FWA in the A-S and A-SW directions, the anterior PM undergoes a displacement that may progressively pull the CTs inserting in the AP commissure, which in turn transfer an increasing load to the commissural region of the leaflets. The auxiliary simulations described in Section 2.6.3, in which the post-FWA annular dynamics was replaced by that of the FTR baseline, yielded systematically higher leaflet stress levels vs. the corresponding simulations with image-based post-FWA annular kinematics (Supplementary Material, Section S1). This result supports the existence of a biomechanically beneficial indirect contribution of FWA on the TA in addition to the direct effect of PM repositioning.

**Fig. 8.**
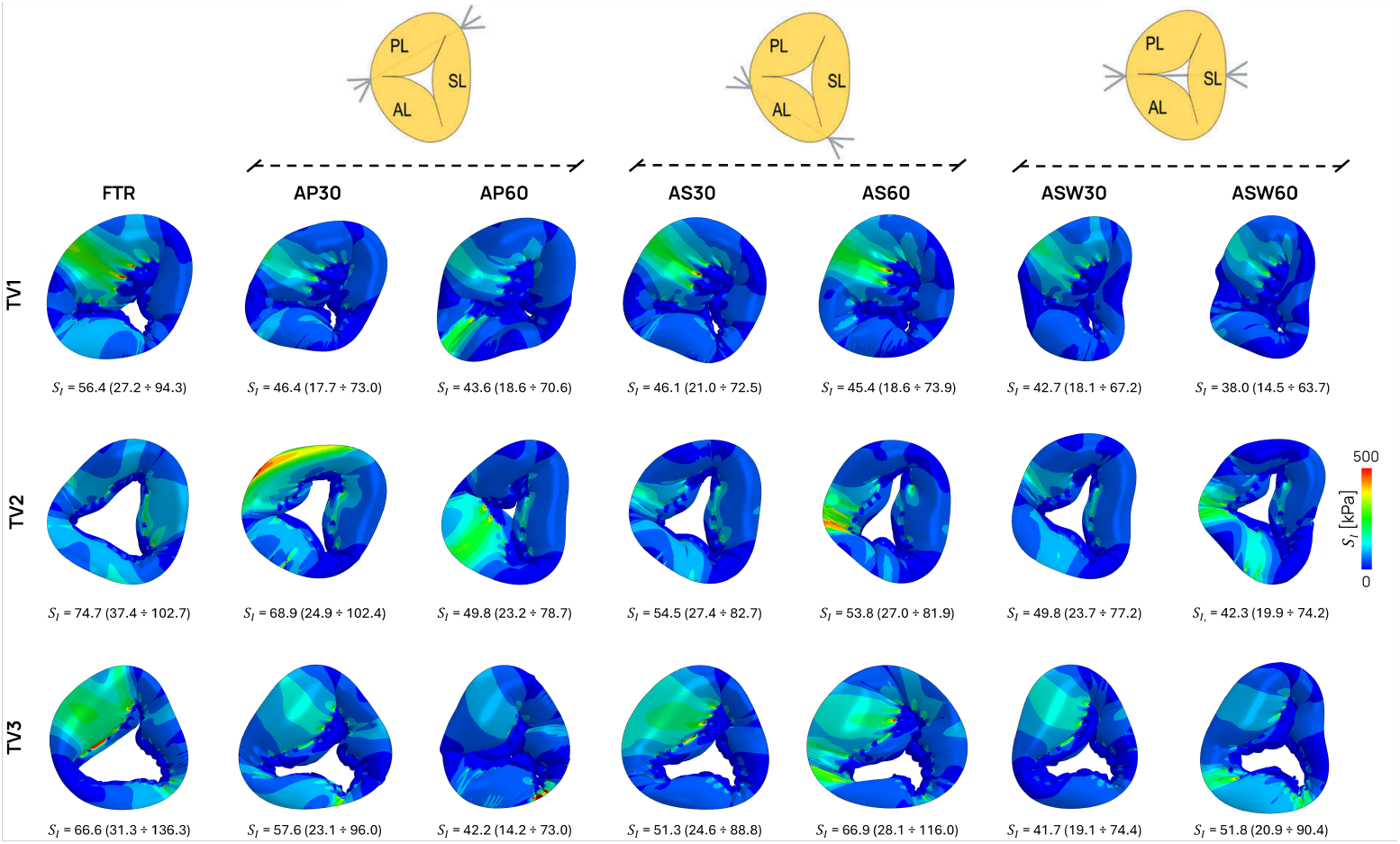
Peak-systolic maximum principal stress (*S*_I_) distributions on the tricuspid valve leaflets under free wall approximation (FWA) conditions for the three analyzed specimens. FWA configurations include anterior-posterior (A-P), anterior-septal (A-S), and anterior-septal wall (A-SW) papillary muscle approximations at 30% and 60% inter-papillary distance reduction levels. Values are reported for each valve as the median with interquartile range (25^th^-75^th^ percentiles) computed over the entire leaflet surface.

### 4.4 Limitations

The present study has several limitations.

First, the FE framework was applied to only three valves, with two of those representing rather extreme FTR conditions. Hence, our findings cannot be generalized, and the framework should be applied to a much larger cohort to verify that, under baseline conditions, the valve configuration at PS can be computed with the level of accuracy reported in the present work, and to explore possible trends in TV systolic strains and stresses with respect to, e.g., the degree of annular dilation or the proportions between different leaflet extent. Likewise, application to a much larger cohort will be necessary to characterize the biomechanical impact of FWA, to verify the trends observed across the different FWA configurations, and to better characterize possible relationships between baseline TV conditions (e.g., degree of malcoaptation and degree of dilation of the underlying RV) and post-FWA systolic biomechanics. Second, leaflet geometries were reconstructed starting from manual segmentation of rt3DE data, which inherently introduces operator-dependent variability and propagates uncertainty into the simulated configurations. Third, leaflet thickness was prescribed parametrically based on literature data [30]. While this assumption may be acceptable in the present setting, where TV models were derived from explanted hearts not subjected to in vivo pathological remodeling, it may become a more relevant limitation when applying the method to imaging acquired in vivo, where leaflet thickening and tissue stiffening associated with chronic FTR are known to occur, leading to deviations from the standard thickness distribution [46].

Moreover, the simulation of FWA itself is affected by two specific limitations. First, PM repositioning was imposed through simplified kinematic assumptions on the distribution of the suture shortening between the two ends of the suture, rather than being computed as the effect of RV wall reshaping induced by FWA and mediated by RV wall mechanical response. This simplification appears reasonable in the moderately regurgitant case (TV1) and along the A-P suture direction, where the simulated configurations matched the in vitro ground truth well, but proved less representative for the A-S and A-SW directions in the more regurgitant valves, where the simulated residual ROA tended to be overestimated. Second, the post-FWA annular dynamics was not predicted by the model but prescribed from post-FWA rt3DE tracings; consequently, the framework cannot be considered fully predictive of the post-operative valve configuration. Overcoming these two limitations would require coupling the TV model with an active model of the RV myocardium, in which PM repositioning would emerge as the mechanical effect of the suture on the ventricular wall and the modification of the TA dynamics would result from the interplay between the FWA and myocardial contraction in the valve plane region. Clearly, such an approach would entail a substantially higher level of complexity, requiring an accurate representation not only of the RV wall anatomy but also of the local myofiber orientation and of the magnitude and timing of myocardial contraction.

## 5 Conclusions

We developed an ultrasound-based FE framework for patient-specific analysis of TV biomechanics in FTR and after FWA-based subvalvular correction. At FTR, simulated leaflet configurations at PS reproduced the ground truth with median geometric errors comparable to the variability of manual segmentation. The preliminary application of the framework to FWA reproduced the in vitro reduction of the ROA and suggested consistent trends across the three analyzed valves, including a progressive reduction of the ROA with the level of approximation and an effect of FWA direction on coaptation restoration. Future integration of the TV model within an active RV myocardial model will enable fully predictive simulations of the post-operative valve configuration.

## Supporting information

s1_section

## Acknowledgements

F.S. gratefully acknowledges the financial support of the “Ricerca Corrente” grant from IRCCS Policlinico San Donato, a clinical research hospital partially funded by the Italian Ministry of Health.

