## Supplementary material for "Rt3DE-based finite element analysis of functional tricuspid regurgitation and RV free wall approximation": s1_section

### S.1 **Effect of post-FWA annular boundary conditions on leaflet stress predictions**

Figure S1 reports the peak-systolic maximum principal stress ($S_I$) distributions on the TV leaflets for TV1, TV2, and TV3 under FTR conditions and after A-P approximation at 30% (AP30) and 60% (AP60), with the latter two simulations repeated using the annular boundary conditions (BCs) of the FTR case (FTR ANNULAR BCs). The comparison was performed to investigate whether the modification of the annular dynamics observed after FWA contributes to the leaflet stress relief predicted by the standard post-FWA simulations. Median $S_I$ values with interquartile range (25th–75th percentiles), in kPa, are reported in Figure S1.

Across all three valves, retaining the FTR annular boundary conditions produced systematically higher median $S_I$ values than the standard post-FWA simulations, with the difference becoming more pronounced at AP60. This supports the existence of a biomechanically indirect effect of FWA on the TA, consistent with the "fourfold mechanism" described by Jaworek et al. [1], and motivates accounting for the post-FWA annular dynamics in the simulation framework.

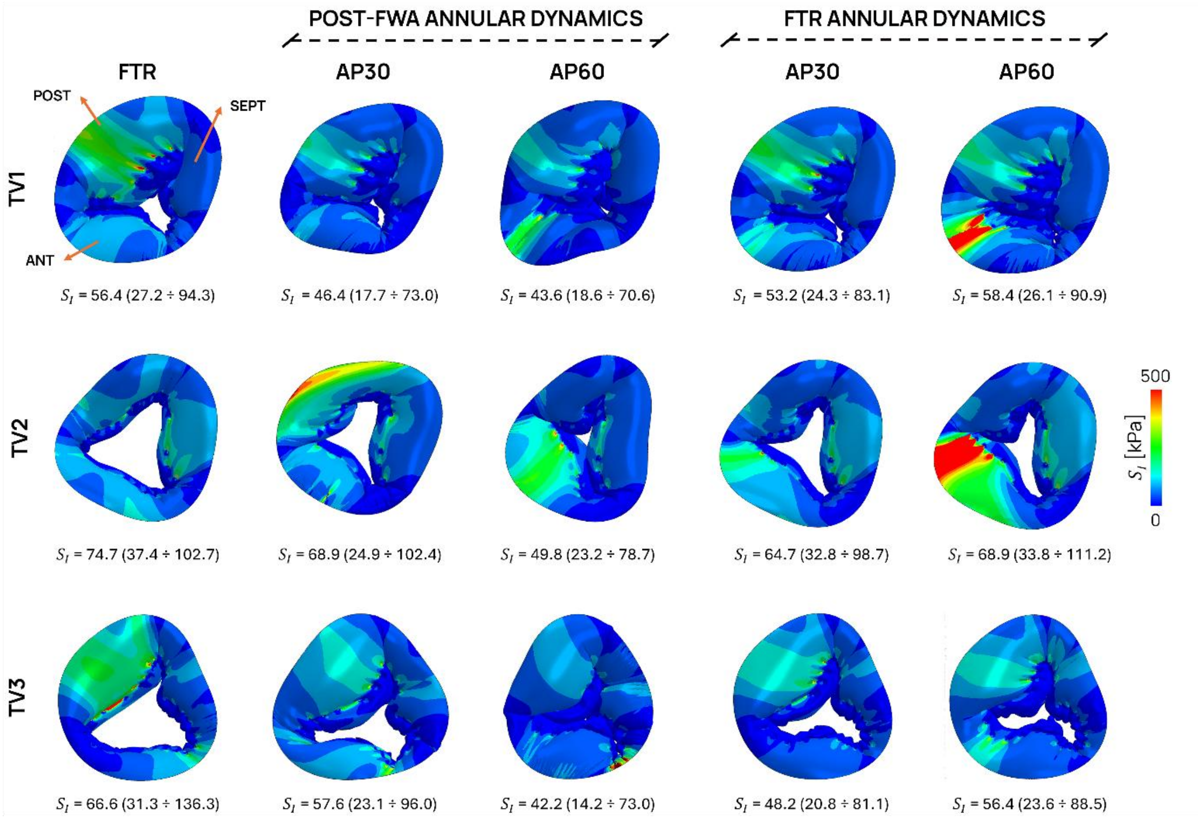


*Figure S1: Peak-systolic maximum principal stress ($S_I$) distributions on the TV leaflets for the three analyzed specimens (TV1, TV2, and TV3) under functional tricuspid regurgitation (FTR) conditions and after anterior-posterior (A-P) free wall approximation at 30% (AP30) and 60% (AP60) inter-papillary distance reduction. For each valve, the two rightmost columns (FTR ANNULAR DYNAMICS) report the corresponding post-FWA simulations performed by replacing the annular boundary conditions derived from post-FWA rt3DE segmentation with those used in the FTR simulation. Values are reported as the median with interquartile range (25th –75th percentiles) computed over the entire leaflet surface.*
